# Population Pharmacokinetics Study of Morinidazole in Patients with Moderate Hepatic Impairment

**DOI:** 10.1101/312728

**Authors:** Yue Kang, Fengyan Xu, Kun Wang, Jing Zhang, Xiaojie Wu, Jufang Wu, Guoying Cao, Jicheng Yu, Beining Guo, Yuancheng Chen, Yingyuan Zhang

## Abstract

**Objective:** Morinidazole is a novel third generation 5-nitroimidazole antimicrobial drug which has demonstrated substantial antibacterial activity against clinical isolates of anaerobe. The aim of this study was to build population pharmacokinetic (PPK) model of morinidazole among patients with hepatic impairment and to provide dosage adjustment strategy for morinidazole in patients with hepatic impairment and/or renal dysfunction.

**Methods:** The nonlinear mixed effects modeling tool NONMEM (version7.3, ICON Development Solutions) was used to develop the PPK model of morinidazole.

**Results:** One-compartment model was conducted to establish the morinidazole PPK model. Disease condition was the significant covariate for CL and weight was the significant covariate for V. The AUC_0-∞_ was 120.44±37.05 (79.25-207.20) μg×h/mL in hepatic impairment group and was 79.46±23.71 (42.94-116.75) μg×h/mL in control group. The AUC_0-∞_ was 164.9±44.8 μg×h/mL and 77.2±23.1 μg×h/mLin in the 3 subjects with both hepatic impairment and mild renal impairment and in the 3 matched healthy subjects, respectively.

**Conclusion:** It is not necessary to adjust morinidazole dosage for patients with moderate hepatic impairment without confirmed renal dysfunction. For patient with moderate hepatic and mild renal impairment, morinidazole regimen should be considered as 500mg every 24 hours. When used in patients with moderate/severe hepatic impairment combined with renal dysfunction, both dosage and interval adjustment of morinidazole should be considered.

## Introduction

Morinidazole is a novel third generation 5-nitroimidazole antimicrobial drug which has demonstrated substantial antibacterial activity against clinical isolates of anaerobe including gram negative *Sporeless bacterium* and gram-positive *coccus* based on the result of pharmacodynamics study *in vitro*. The antibacterial activity of morinidazole against *Bacteroides fragilis*, *Veillonella* and *Clostridium perfringens* is comparable to that of ornidazole, which is 2 to 8 times stronger than that of metronidazole and tinidazole. The antibacterial activity of morinidazole against *Bacteroides distasonis* and *Bacteroides ovatus* is comparable to that of ornidazole, which is 2 to 8 times stronger than that of metronidazole and tinidazole as well (1, 2). Morinidazole and Sodium Chloride Injection was approved by CFDA in February 2014 for the treatment of pelvic inflammatory disease caused by *Peptostreptococcus*, *Bacteroides fragilis, Veillonella* and *Bacteroides distasonis*, suppurative appendicitis and gangrenous appendicitis caused by anaerobes(3).

Morinidazole showed a positively correlated relationship between dosage and AUC_0-t_ as well as C_max_. The V_ss_ of morinidazole was 1209±158 mL/kg after infusion at 16mg/kg for 2h. The human plasma protein binding rate of morinidazole was 22.1 to 27.2%. Morinidazole was widely distributed in tissues and body fluids. Morinidazole was mainly metabolized by glucuronidation (mediated by UGT1A9 enzyme) and sulfation rather than CYP450 in human. It took 36h for 70% morinidazole to excrete as unchanged drug and phase II metabolites via renal pathway in healthy subjects(3).

Phase I to phase III clinical trials had been completed before morinidazole launching. Our team had finished the pharmacokinetics study of morinidazole in moderate hepatic impaired subjects in 2010, the results showed the AUC_0-∞_ of hepatic impaired subjects and healthy subjects is 116.3±41.8 μg×h/mL and 77.2±25.3 μg×h/mL, respectively. The AUC_0-∞_ of 3 hepatic impaired subjects with mild renal function impaired was 166.7±52.1 μg×h/mL. The metabolism rate of a nitrogen-oxide (N-oxide), M2, was low (2). Meanwhile, the pharmacokinetics study of morinidazole among severe renal function impaired subjects had been conducted. The AUC of severe renal function impaired subjects was 1.4 times higher than that of healthy subjects(4). However, non-compartment model analysis which published before cannot assess the impact of patient physiology and pathology condition on pharmacokinetics(PK) parameters. Hence, compartment model was performed in order to assess the impact of variables on AUC in this study. In clinical practice, some patients who experiencing infections have complications of liver functions impairment and/or renal function impairment, while some elderly patients have liver function impaired with physiologically decreased glomerular filtration rate. Moreover, the PK study among hepatic function impaired patients used creatinine clearance rate (CCr) to assess the renal function of subjects. The guideline of KDIGO recommends assessing renal function by estimated glomerular filtration rate (eGFR)(5). The value is always not equal between CCr and eGFR(6), thus the renal function was reassessed in this study. The aim of this study was to build the PPK model of morinidazole among liver dysfunction patients and explore the covariate in hepatic impairment patient as well as to compare the PK parameter changes in patients with liver dysfunction combined with renal dysfunction by model simulation in order to provide the dosage adjustment strategy of morinidazole in the special populations and to guide the rational use.

## Methods

### 1. Data of Study for PPK Modeling

Data from the PK study of morinidazole among patients with hepatic function impairment (Study No. ChiCTR1800015771, approved by HIRB of Huashan hospital, Fudan University on 6^th^ November 2009) (2), including demographic characteristics, laboratory test results and plasma concentrations were used for PPK analysis. The study was an open-label, single dose, parallel control study in which 12 subjects were enrolled in the moderate liver function impaired group and the healthy controlled group, respectively. Single dose of 500mg morinidazole and sodium chloride injection (500mg/100ml) was administrated via intravenous infusion for 45min. All the blood samples were collected from pre-dosing to within 48h after infusion.

#### Subjects

Individuals with moderate (Child-Pugh class B) hepatic impairment caused by hepatitis B induced cirrhosis were enrolled in hepatic impaired group. The healthy control subjects were matched based on the age (±5 years old), gender and weight (±15%) of liver function impaired subjects. CKD-EPI equation (2009) was used to reevaluate the eGFR of the subjects(7). Moreover, model for end-stage liver disease (MELD) and Albumin-Bilirubin (ALBI) score were performed to reassess the liver function of subjects from liver function impaired group. Demographic characteristics were shown in Table 1

**Table 1.** Demographic Characteristics of subjects in the PK study among moderate liver dysfunction population

|  | Healthy subject Group<br>(N=12) | Patient Group<br>(N=12) | Total<br>(N=24) |
| --- | --- | --- | --- |
| Gender, male (%) | 7 (58.33) | 7 (58.33) | 14 (58.33) |
| Age(years) | 47.00 (35.00-67.00) | 51.00 (37.00-65.00) | 48.50<br>(35.00-67.00) |
| Weight(kilograms) | 60.50 (47.00-78.50) | 66.50 (44.50-92.00) | 63.00<br>(44.50-92.00) |
| Height(centimeters) | 161.00<br>(150.00-179.00) | 169.50<br>(155.00-180.00) | 165.50<br>(150.00-180.00) |
| BMI(kg/m<sup>2</sup>) | 23.98 (19.27-27.82) | 23.61 (17.83-28.40) | 23.72<br>(17.83-28.40) |
| **Hepatic Function** |  |  |  |
| ALT(U/L) | 16.00 (11.00-53.00) | 40.00 (22.00-89.00) | 23.50<br>(11.00-89.00) |
| AST(U/L) | 20.50 (12.00-28.00) | 59.50<br>(24.00-175.00) | 27.50<br>(12.00-175.00) |
| **Child-Pugh** |  |  |  |
| Score, n (%) |  |  |  |
| Normal | 12 (100.00) | 0 | 12 (50.00) |
| 7 | 0 | 5 (41.67) | 5 (20.83) |
| 8 | 0 | 2 (16.67) | 2 (8.33) |
| 9 | 0 | 5 (41.67) | 5 (20.83) |
| **Renal Function** |  |  |  |
| Cr(μmol/L) | 62.00 (45.00-85.00) | 61.50<br>(36.00-101.00) | 61.50<br>(36.00-101.00) |
| eGFR(ml/min/1.73m<sup>2</sup>) | 105.00<br>(90.00-120.00) | 106.50<br>(70.00-122.00) | 105.00<br>(70.00-122.00) |
| **Concomitant Medication,** |  |  |  |
| antiviral agent, n (%) | 0 | 12 (100) | 12 (50) |

BMI: body mass index; ALT: alanine aminotransferase; AST: aspartate transaminase;
Cr: creatinine; eGFR: estimated glomerular filtration rate

#### Study Design

Plasma samples were collected prior to administration of morinidazole, 22.5min after start of the infusion and at 0h, 0.25, 1, 2, 4, 6, 8, 12, 24, 36, 48h after the end of infusion. Liquid chromatography-tandem mass spectrometry (LC-MS/MS) was performed to determine the concentration of unchanged morinidazole.

#### Safety Assessment

Blood samples were collected to assess the liver and renal function in screening period, on the dosing day, as well as at 24h and 48h after administration of morinidazole. The safety assessment included results analysis of clinical manifestations, vital signs, physical examination, laboratory tests and 12-lead electrocardiogram (ECG).

#### Data description

A total of 336 concentrations samples from 24 subjects were used to establish the PPK model.

### 2. PPK Modeling Building

#### Basic Modeling Building

Tow-compartment and one-compartment model was evaluated to describe the pharmacokinetics of morinidazole indications. A modeling analysis was conducted using the nonlinear mixed effects modeling tool NONMEM (version7.3, ICON Development Solutions).

#### Covariate analysis

The model was based on the first-order conditional estimation (FOCE) method with η-ε interaction. Model selection was based on likelihood ratio tests, residual analysis and parameter rationalities. Covariate analysis was conducted after the selection of the base model. Scatter plots of parameter estimates from the selected base model and potential covariates (including: disease condition (moderate hepatic impairment or health), gender, height, weight, body mass index(BMI), alanine aminotransferase(ALT), aspartate transaminase(AST), ascites, total bilirubin(TB), Child-Pugh score and estimated glomerular filtration rate (eGFR) were plotted to explore covariate-parameter relationships. The Stepwise Covariate Model (SCM) module in Perl-speaks-NONMEM software (version 3.4.2, Uppsala University, Sweden) was then used for covariate screening and identification.

Covariate relationships as linear function, exponential function and power function were assessed for continuous covariates, and a stepwise approach was used to evaluate covariate effects. The statistical criteria for a covariate to be incorporated in the model were a decrease of >3.84 in the OFV (p=0.05, χ^2^ distribution with one degree of freedom) in the forward step and an increase of <6.63 in the OFV (p=0.01, χ^2^ distribution with one degree of freedom) in the backward step. A decrease in objective function value (OFV) >6.63 was considered statistically significant.

#### Model assessment and validation

The goodness-of-fit was evaluated by comparing the following graphs both in base model and in final model: observed concentrations versus population predictions (DV vs PRED), observed concentrations versus individual predictions (DV vs IPRE), conditional weighted residuals (CWRES) versus time, and CWRES vs PRED.

A bootstrap resampling technique was used for the model validation. The advantage is that calculation of certain statistics with a nonparametric bootstrap method does not depend on the assumption of sample distribution. The basic idea is to use the Monte Carlo method repeated sampling with replacement in the sample, thus forming a self-sampling. Once the self-sampling reaches a certain frequency, it forms a statistical distribution, and the original sample statistics can then be estimated with a semi-empirical method. Visual predictive check (VPC) was used to graphically assess the appropriateness of the compartment model because of the importance of the ability of the model to simulate data similar to the original data. The concentration profiles were simulated 1000 times and compared with observed data to evaluate the predictive performance of the model.

#### Post-hoc analysis and Simulations

The AUC_0-24_ of individuals and the change of AUC_0-24_ based on Child-Pugh score was simulated by PK parameters and Intra-and inter-individual variation based on the final model

#### Individual Parameters

Descriptive statistical analysis was used to analyze the PK parameters of 24 subjects which calculated by population pharmacokinetics method by NONMEM via SPSS (Version 17.0). Meanwhile, the hepatic function of the 12 liver impaired subjects was reassessed by MELD and ALBI score to observe the correlation between MELD /ALBI score and AUC of morinidazole.

#### Correlation Analysis Between Covariate and Morinidazole AUC_0-24_

Spearman’s correlation was conducted by SPSS (Version 17.0) to analyze the correlation between the covariates and morinidazole AUC_0-24_. The covariates included ALT, AST, GGT, TB, ABL, prolongation of the prothrombin time and eGFR.

## Results

### 1. PPK Modeling Analysis

#### Basic Model

The plasma concentration of 24 subjects was used to build the model (Figure 1). The Child-Pugh score of 12 moderate hepatic impaired subjects caused by chronic virus hepatitis induced cirrhosis was from 7 to 9 (average was 8). Both one-compartment and two-compartment model were performed to establish the basic model. One-compartment model fit the data more precisely than two-compartment model and thus was selected as the basic model. The objective formulation value (OFV) was 4502.814. The parameters of PPK included CL (5.32 L/h) and V_c_ (50.5 L) of morinidazole. The inter-individuals variation was well descripted by the index model.

**Figure 1.**
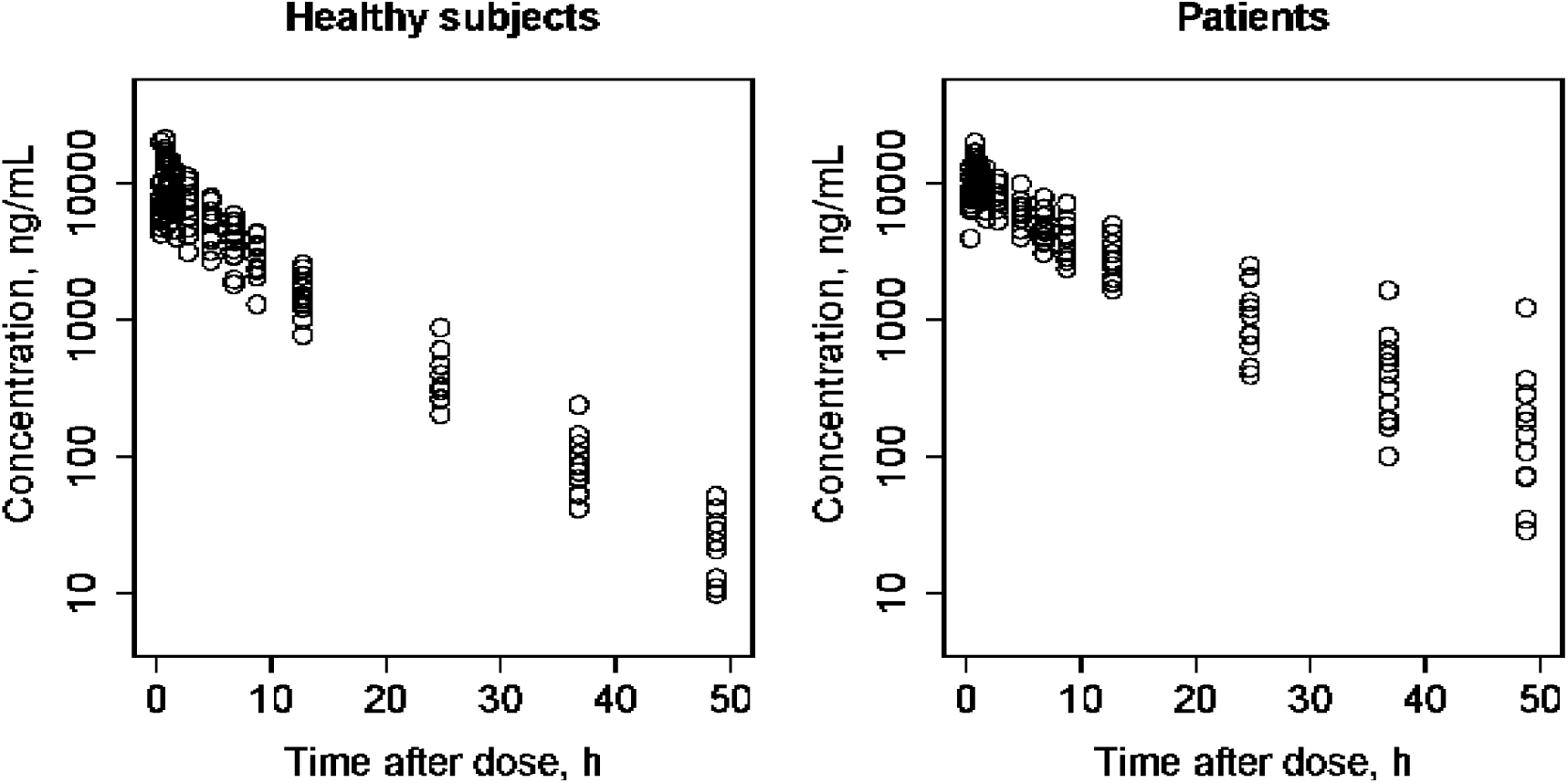
Unchanged Morinidazole Plasma Concentration Data for model building.

#### Covariate Analysis

Scatter plots of parameter estimates from the selected basic model and potential covariates showed that height, weight, disease condition, hepatic encephalopathy, prolongation of the prothrombin time and ALB (P<0.01) as well as moderate hepatic impairment combined with eGFR ⩽ 90ml/min/1.73m^2^, ascites, TB and Child-Pugh score (0.01<P<0.05) were the potential covariates. As a result, disease condition, ALB, Child-Pugh score, ascites, TB and eGFR were selected as potential covariates for CL. Weight was chose as the potential covariate for V. It was showed that disease condition was the significant covariate for CL and weight was the significant covariate for V via forward selection and backward deletion. The covariate equation was as the following:

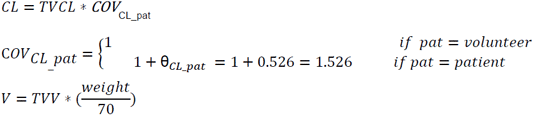

θ_*CL–pat*_is the changed value of dichotomy covariate of COV_cl_ in moderate hepatic impairment patients.

#### Modeling evaluation and validation

The population parameter estimates of the final model for CL and V were illustrated in Table 2. The population estimate values (RSE%) were 4.32 L/h (8.30%) for CL and 55.5 L (4.80 %) for V. Incorporation of patient status and weight as covariate effects on CL and V decreased the OFV from 4502.814 to 4483.082.

**Table 2.** the result of PK parameters by 1000 bootstrap via final model

|  | Estimate(RSE%) | 1000 successful bootstrap median (95% CI) |
| --- | --- | --- |
| PK Parameter |  |  |
| CL (L/h) | 4.32 (8.30) | 4.32 (3.64-5.13) |
| V (L) | 55.5 (4.80) | 55.4 (50.8-60.8) |
| $\theta_{CL\_pat}$ | 0.521 (35.5) | 0.524 (0.185-0.943) |
| Interindividual variability |  |  |
| CL, % | 28.9 (13.6) | 27.2 (18.3-34.8) |
| V, % | 21.3 (19.9) | 20.5 (11.5-28.1) |
| Residual error |  |  |
| Prop, % | 21.8 (4.80) | 21.9 (19.8-24.2) |

CI: Confidential interval

The GOF figure of model evaluation showed that both population and individual predictions were consistent with observation value and the final model fit better than the basic model. The figure of Individual prediction vs. observation, Population prediction vs. observation, Conditional weighted residual vs population prediction and Conditional weighted residual vs time was in Figure 2

**Figure 2.**
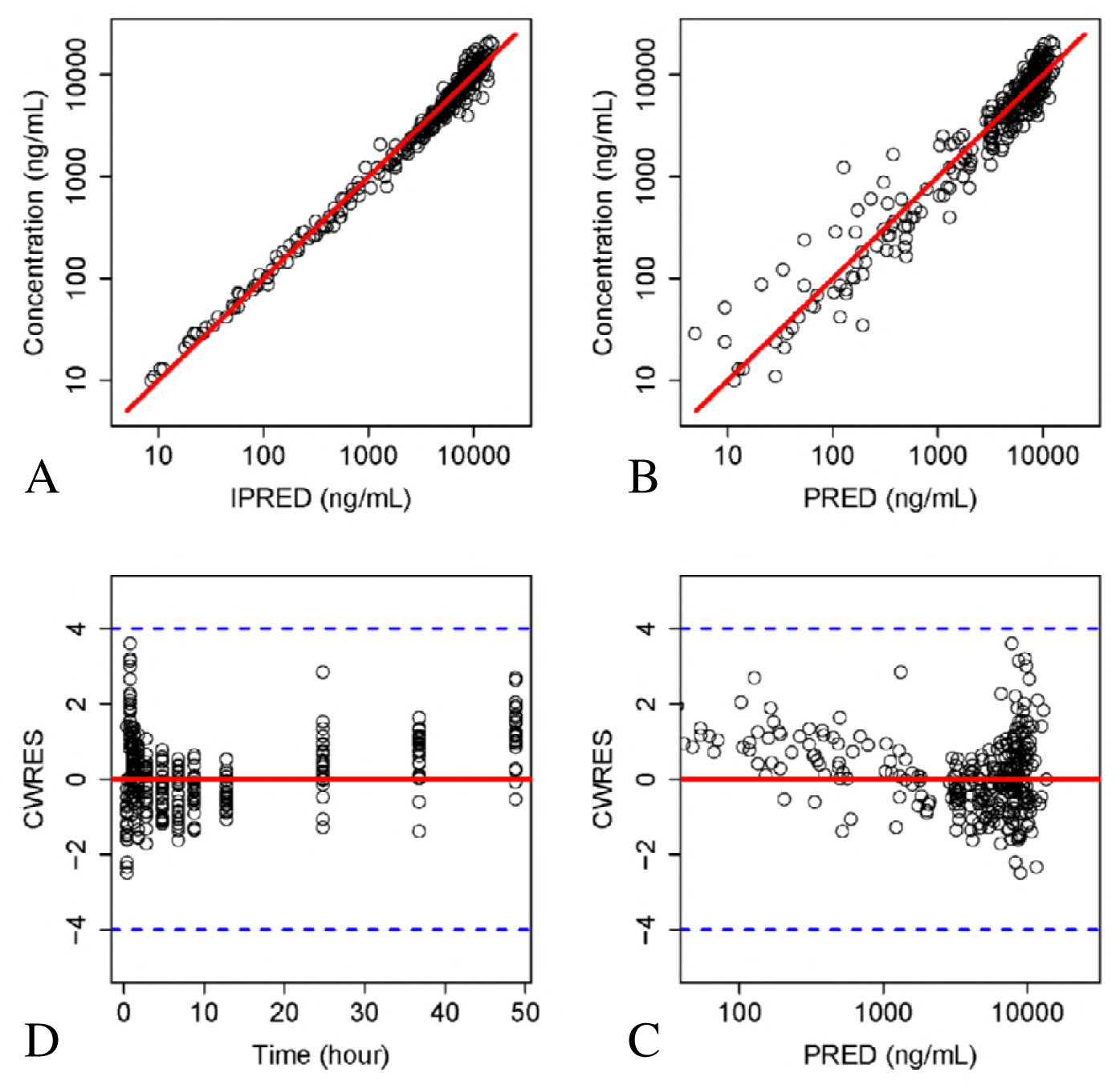
General goodness-of-fit plots for PPK final model. A)Individual prediction vs. observation; (B) Population prediction vs. observation (C)Conditional weighted residual vs population prediction; (D) Conditional weighted residual vs time; The red line in (A) and (B) represent regression line, respectively, whereas in (C) and (D) are the position where conditional weighted residual equal 0.

In Figure 2, the red line in (A) and (B) represented regression line, respectively, whereas in (C) and (D) were the position where conditional weighted residual equal 0.

The final model was validated by nonparametric bootstrap. The results demonstrated a complete overlap of the 95% confidence interval of parameter estimates with the ranges of the 2.5^th^ to 97.5^th^ percentiles, indicating that the final model was robust (Table 2). The final model demonstrated strong stability, with 1000 bootstrap runs fitting successfully, and bootstrap estimates very similar to the population estimates.

Figure 3 showed the results of VPC using 1000 monte carlo simulations. The observed concentration was included in the range of confidence intervals and the median and 95% confidence interval lines were located near the middle area of the 1000 results, suggesting the sufficiency of the predictive power of the model.

**Figure 3.**
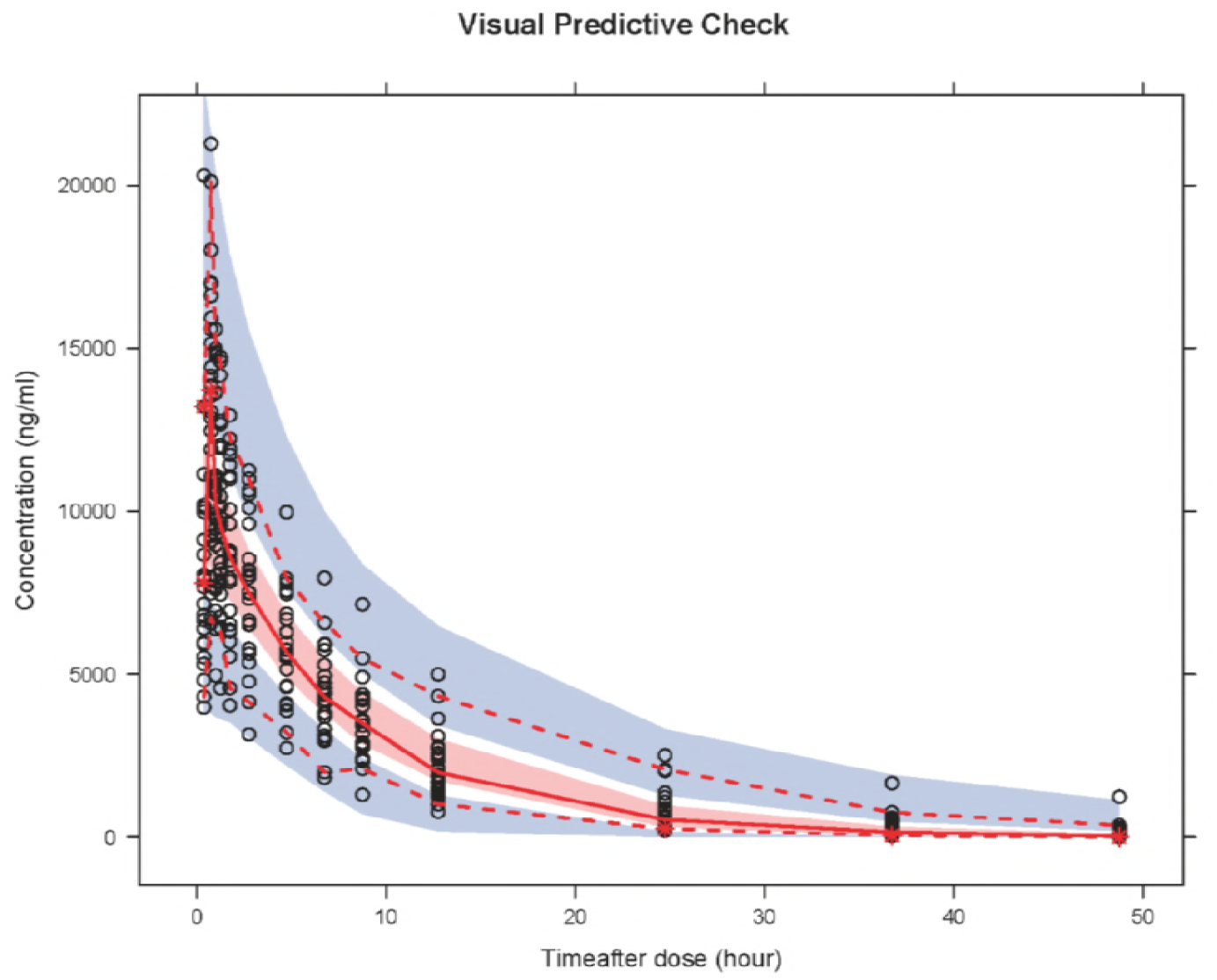
VPC results validated the prediction ability of final model. Open circles represent observed concentrations, the solid line and the dashed line represents the median and the 95*%* CI of observation, respectively. The middle red shadow areas represent the 95*%* confidence intervals of median for the results of 1000 times simulation of final model and the blue shadow areas represent the 95*%* confidence intervals of the 2.5th and 97.5th percentiles of the results of 1000 times simulation of pharmacokinetic final model.

Open circles represented observed concentrations. The solid line and the dashed line represented the median and the 95% CI of observation, respectively. The middle red shadow areas represented the 95% confidence intervals of median for the results of 1000 times simulation of final model and the blue shadow areas represented the 95% confidence intervals of the 2.5th and 97.5th percentiles of the results of 1000 times simulation of pharmacokinetic final model.

#### PK Parameters of Individuals

The AUC_0-∞_ was 120.44±37.05 (79.25-207.20) μg×h/mL in hepatic impairment group and was 79.46±23.71 (42.94-116.75) μg×h/mL in control group. Other PK parameters were listed in table 3. There were 3 subjects from hepatic impairment group combined with eGFR ⩽90ml/min/1.73m^2^. Although the eGFR of one subjects was 90 ml/min/1.73m^2^, the urine routine test revealed weakly positive urine protein and occult blood. Therefore, the subject was diagnosed with mild renal function impairment. The AUC_0-∞_, C_max_, CL, V and t_1/2_ of the three subjects were 164.9±44.8 (range was from 117.9 to 207.2) μg×h/mL, 16.53±3.62μg/mL, 3.2±0.94L/h, 44.65±7.9L and 10.18±3.45h, respectively. While the AUC_0-∞_, C_max_, CL, V and t_1/2_ of the 3 matched healthy subjects were 77.2±23.1 (range was from 54.9 to 101.0) μg×h/mL, 12.13±4.61μg/mL, 6.89±2.09L/h, 54.04±19.98L and 5.39±0.49h respectively. The PK parameters of moderate hepatic impairment subjects without renal function impairment and the controlled healthy subjects were listed in table 3. The correlation between AUC_0-24_ and Child-Pugh score was showed in Figure 4.

**Table 3.** Descriptive Analysis of Individuals PK Parameters by Different Groups

| PK Parameters | Subjects of moderate liver function impaired combined with renal function imparied N=3 | The matched healthy subjects N=3 | Subjects of moderate liver function impaired with normal renal function N=9 | The matched healthy subjects N=9 | Subjects of moderate liver function impaired N=12 | The matched healthy subjects N=12 |
| --- | --- | --- | --- | --- | --- | --- |
| CL (L/h) |  |  |  |  |  |  |
| Mean±SD, | 3.20±0.94 | 6.89±2.09 | 4.89±0.97, | 6.85±2.31 | 4.47±1.19 | 6.86±2.17 |
| (Range) | (2.41-4.24) | (4.95-4.16) | (3.85-6.31) | (4.28-11.64) | (2.41-6.31) | (4.28-11.64) |
| CI90% | 1.62-4.78 | 3.36-10.42 | 4.30-5.49 | 5.41-8.28 | 3.85-5.09 | 5.73-7.98 |
| V (L) |  |  |  |  |  |  |
| Mean±SD, | 44.65±7.90 | 54.04±19.98 | 52.05±8.51, | 54.20±19.11 | 50.20±8.67 | 54.16±18.39 |
| (Range) | (35.53-49.30) | (39.04-76.72) | (40.20-66.1) | (31.68-88.02) | (35.53-66.09) | (31.68-88.02) |
| CI90% | 31.33-57.98 | 20.36-87.73 | 46.77-57.32 | 42.35-66.05 | 45.70-54.69 | 44.62-63.69 |
| C<sub>max</sub> (mg /L) |  |  |  |  |  |  |
| Mean±SD, | 16.53±3.62 | 12.13±4.61 | 13.38±2.39 | 13.05±5.18 | 14.17±2.93 | 12.82±4.85 |
| (Range) | (12.87-20.11) | (6.82-15.15) | (9.24-17.01) | (6.94-21.28) | (9.24-20.11) | (6.82-21.28) |
| CI90% | 10.42-22.63 | 4.36-19.90 | 11.90-14.86 | 9.85-16.26 | 12.65-15.69 | 10.30-15.34 |
| t<sub>1/2</sub> (h) |  |  |  |  |  |  |
| Mean±SD, | 10.18±3.45 | 5.39±0.49 | 7.55±1.58 | 5.48±0.59 | 8.21±2.32 | 5.45±0.55 |
| (Range) | (8.03-14.16) | (4.86-5.83) | (5.37-10.23) | (4.67-6.18) | (5.37-14.16) | (4.67-6.18) |
| CI90% | 4.36-15.99 | 4.56-6.21 | 6.27-8.53 | 5.11-5.84 | 7.00-9.41 | 5.17-5.74 |
| AUC<sub>0-∞</sub> |  |  |  |  |  |  |

| (µg×h/mL) |  |  |  |  |  |  |
| --- | --- | --- | --- | --- | --- | --- |
| **Mean±SD,** | 164.91±44.84 | 77.18±23.09 | 105.62±19.92 | 80.22±25.24 | 120.44±37.05 | 79.46±23.71 |
| **(Range)** | (117.90-207.20) | (54.87-100.98) | (79.25-129.78) | (42.94-116.75) | (79.25-207.20) | (42.94-116.75) |
| **CI90%** | 89.32-240.50 | 38.24-116.11 | 93.28-117.97 | 64.58-95.87 | 101.24-139.65 | 67.17-91.75 |

| C<sub>24h</sub> (ug/ml) |  |  |  |  |  |  |
| --- | --- | --- | --- | --- | --- | --- |
| **Mean±SD,** | 2.11±0.94 | 0.45±0.14 | 1.06±0.43 | 0.48±0.18 | 1.32±0. 72 | 0.47±0.17 |
| **(Range)** | (1.28-3.13) | (0.35-0.61) | (0.56-1.67) | (0.24-0.86) | (0.56-3.13) | (0.24-0.86) |
| **CI90%** | 0.527-3.697 | 0.207-0.687 | 0.797-1.328 | 0.37-0.60 | 0. 951-1.698 | 0.385-0.561 |

| AUC<sub>0-24</sub> |  |  |  |  |  |  |
| --- | --- | --- | --- | --- | --- | --- |
| (µg×h/mL) |  |  |  |  |  |  |
| **Mean±SD,** | 130.90±24.18 | 73.69±22.24 | 93.23±14.47 | 76.34±24.14 | 102.65±23.43 | 75.68±22.70 |
| **(Range)** | (103.04-146.47) | (51.70-96.17) | (74.05-109.34) | (41.15-112.20) | (74.05-146.47) | (41.15-112.20) |
| **CI90%** | 90.14-171.67 | 36.20-111.18 | 84.26-102.20 | 61.38-91.30 | 90.50-114.79 | 63.91-87.45 |

**Figure 4.**
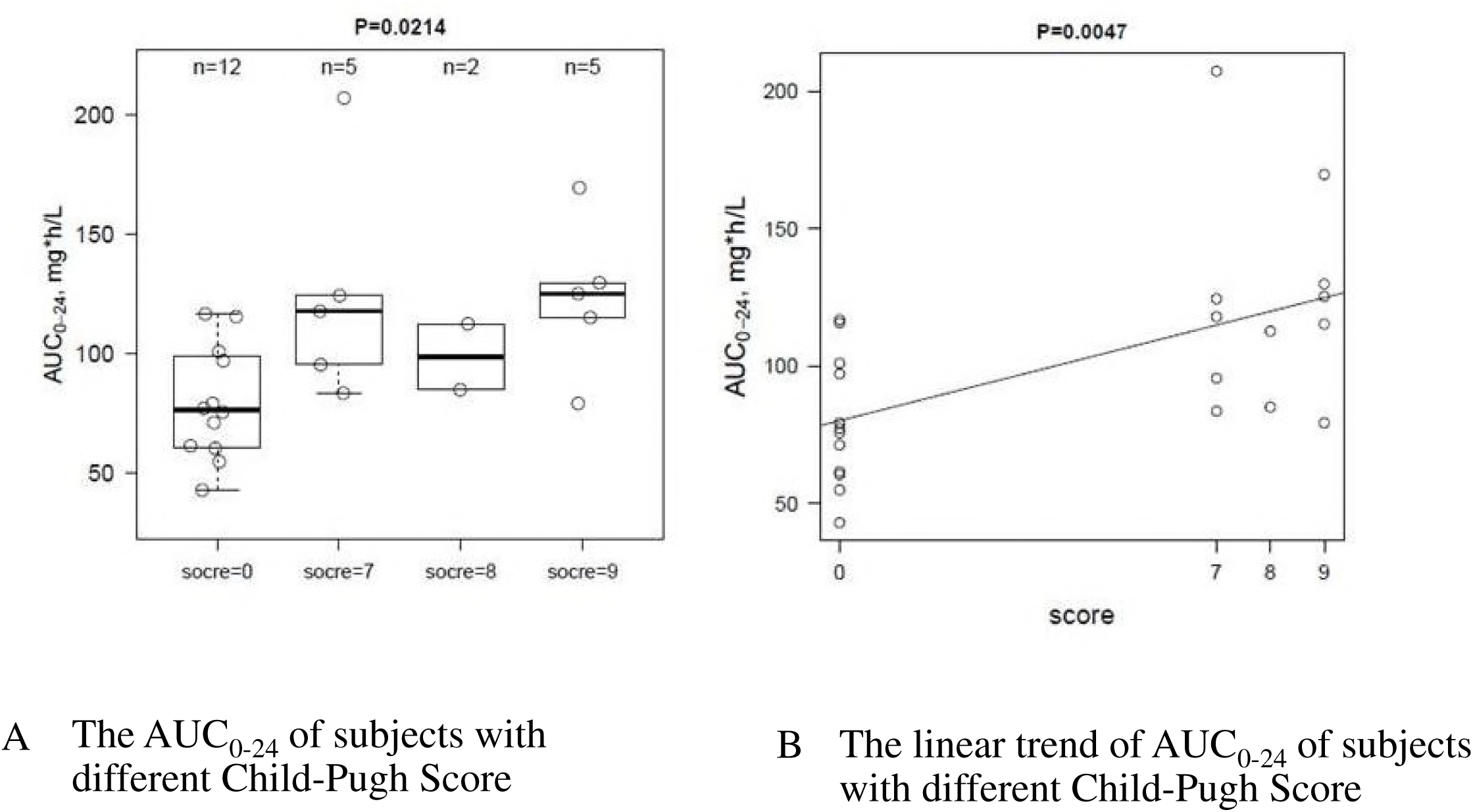
the correlation between AUC0-24 and Child-Pugh score.

MELD score and ALBI score were used to reassess the liver function of subjects in hepatic impaired group. The results were showed in table 4 and figure 5.

**Figure 5.**
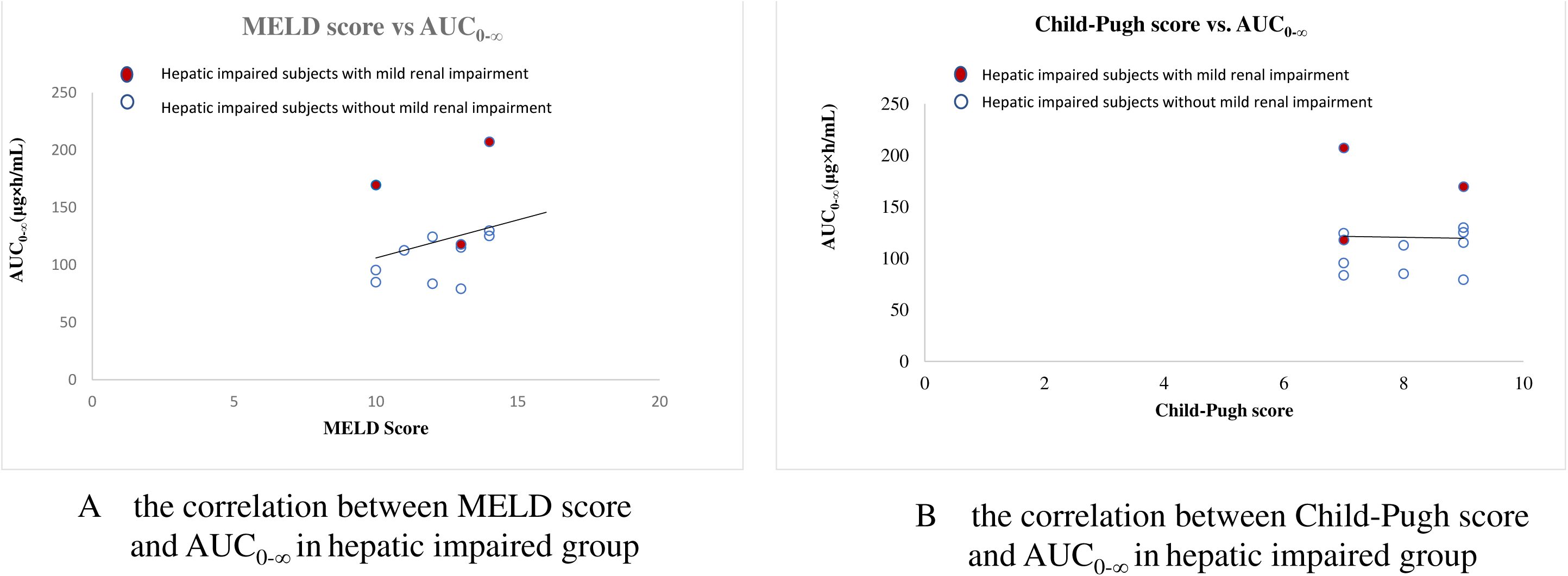
The linear trend of AUC0_-∞_ of subjects with different MELD and Child-Pugh Score.

**Table 4.** MELD score and ALBI score of subjects in hepatic impaired group

| No. | Child-Pugh<br>score | MELD<br>score | ALBI<br>score | ALBI<br>grade | eGFR(ml/mi<br>n/1.73m2) | AUC<sub>0-∞</sub><br>(μg×h/mL) |
| --- | --- | --- | --- | --- | --- | --- |
| 1 | 9 | 10 | -1.36449 | 3 | 90 | 169.618 |
| 2 | 9 | 13 | -1.98412 | 2 | 100 | 79.25437 |
| 3 | 8 | 10 | -2.26514 | 2 | 121 | 85.0051 |
| 4 | 7 | 13 | -1.3875 | 3 | 72 | 117.8995 |
| 5 | 9 | 13 | -1.93767 | 2 | 102 | 115.2206 |
| 6 | 7 | 14 | -1.69243 | 2 | 70 | 207.2024 |
| 7 | 9 | 14 | -1.92343 | 2 | 96 | 129.7757 |
| 8 | 7 | 12 | -1.9492 | 2 | 111 | 124.4122 |
| 9 | 7 | 12 | -1.88932 | 2 | 120 | 83.55615 |
| 10 | 7 | 10 | -1.48837 | 2 | 119 | 95.50186 |
| 11 | 8 | 11 | -1.79168 | 2 | 112 | 112.6202 |
| 12 | 9 | 14 | -1.67024 | 2 | 122 | 125.2662 |

#### Spearman’s rank correlation coefficient analysis

Spearman’s rank correlation coefficient was used to analyze the correlation between liver function index, Child-Pugh index, eGFR and AUC_0-24_. The result showed that the correlation coefficient of AST, ALT and PT was less than 0.05.

### Discussion

To our knowledge, this study was the first study to explore the population-pharmacokinetics of morinidazole injection. Both one-compartment and two-compartment model were performed in the study. The result showed that one-compartment model with first-order elimination was optimal for pharmacokinetic data modeling.

Moderate haptic impairment was the significant covariate for CL and the weight was the significant covariate for V based on the result of this PPK study.

Hepatic impairment is mainly caused by virus hepatitis induced cirrhosis in China(8). Hepatitis B was the cause of hepatic impairment of all the 12 subjects. Child-Pugh score is recommended by FDA to assess the liver function in the PK study of hepatic impaired population(9). Besides Child-Pugh score, MELD score and ALBI Grade are conducted to assess the liver function in clinical practice as well. Child-Pugh is widely used with the index of the synthetic function (ALB and PT) and the elimination function (TB) of liver. It is known that renal function, affecting the PK of morinidazole, is also an important index to predict the prognosis of hepatic function whilst not included in Child-Pugh score. (10) Therefore, the other two score were used to reassess the liver function of the subjects in this study.

MELD score uses serum creatinine, international normalized ratio (INR) and TB as indexes, which reflect the renal function, synthetic and elimination function of liver. The equation of MELD is 3.78 × Ln(TB μmol**/**L) + 11.2 × Ln(INR) + 9.57 × Ln(creatinine mg**/**dL) + 6.4(11) (10). ALBI grade demonstrated an optimal assessment method to predict the prognosis and mortality of liver function impaired which is also in advantage of treatment choice demonstrated by some recent studies. Only TB and ALB are used as covariates in the grade assessment of severity of hepatic impairment(12-14). MELD score showed positive correlation with the change of morinidazole AUC_0-∞_ in our study probably due to the covariate of serum creatinine in the formula. However, the sample size was not adequate to analyze the statistical significance of the correlation. Hence a further study is needed to analyze which liver function assessment method is the best to predict the change in morinidazole PK parameters in hepatic impaired population.

Pearman’s rank correlation coefficient analysis showed that TB and eGFR had no significant correlation with AUC_0-24_, but PT, ALB and AST had significant correlation with AUC_0-24_. However, ALT and AST are the markers of hepatocellular damage rather than liver function(15). A small sample size of only 3 patients with impaired renal function could be the reason why there is no significant correlation between eGFR and AUC_0-24_. The other reason could be that single factor index is not optimal to predict morinidazole PK parameters. No correlation between TB an AUC_0-24_ was probably attributed to the major renal elimination pathway of morinidazole(4). Therefore, the level of PT and ALB could probably be the prediction index for the morinidazole AUC change among hepatic impaired population, but it needs further studies to validate.

A PK study of a single dose of 500mg morinidazole in healthy subjects reported that the C_max_ was 10.8±1.88μg/ml, AUC_0-∞_was 72.5±14.5 μg×h/mL and t_1/2_ was 5.75±0.66h(3).In Zhang’s PK study of morinidazole in severe among renal function impaired population showed that after administrating a single dose of morinidazole the AUC_0-∞_, CL, C_max_ and t_1/2_ of controlled healthy subjects were 71.94±15.95 μg×h/mL, 7.21±1.64L/h, 13.74±3.14μg/mL and 5.54±0.63h, respectively, while the AUC_0-∞_, CL, C_max_ and t_1/2_ of severe renal impaired subjects were 102.79±15.95 μg×h/mL, 5.21±1.47L/h, 11.65±2.97μg/mL and 7.52±1.17h respectively. The AUC_0-∞_ of severe renal impaired subjects was 1.4 time higher than that of the controlled healthy subjects. Although the major metabolite (M4-1) of morinidazole was 7 times higher in severe renal impaired subjects, the concentration was low(4). So the dosage adjustment strategy was referred to PK parameters of unchanged morinidazole, not the metabolite.

Our PPK study showed that the ratio of AUC_0-∞_ of moderate hepatic impaired subjects without impaired renal function to AUC_0-∞_ of healthy matched subjects was less than 2, meanwhile no SAE was reported from both groups in the PK study. Therefore, there is no need to adjust the dosage of morindazole in patients with moderate hepatic impairment and normal renal function. However, since the AEs incidence had significant difference between the two group with a higher incidence in liver function impaired group(2), safety monitoring during the administration of morinidazole was considered for the hepatic impaired group. The ratio of AUC_0-∞_ of moderate hepatic impaired subjects combined with mild renal function impairment to AUC_0-∞_ of healthy matched subjects was more than 2. As a result, dosage adjustment should be considered. The PK study of morinidazole in healthy subjects showed a positively correlated relationship between dosage and AUC_0-t_ (r was 0.979) (3). Morinidazole showed a linear relationship between dosage and AUC. Morinidazole is a concentration-dependence antimicrobial agent and the PK/PD index is AUC/MIC(16). The t_1/2_ of moderate liver function impaired subjects with mild renal function impairment was prolonged by 1.86 times than that of healthy controlled subjects. Therefore, a prolonged administration interval without dose adjustment of 500mg per day should be considered.

Dosage adjustment by both dosage and interval should also be considered among the population combined with both moderate liver function impaired and moderate/severe renal function impaired or both severe impaired liver function and renal function. The administration adjustment strategy requires a further study. Whether the dosage adjustment among both renal and mild liver dysfunction patients also needs further data collections from clinical practice.

Elderly always combined with physiologically decreased glomerular filtration rate(GFR). One study showed that there was 68% elderly (73.7±6 year) who had the creatinine clearance rate (CCr) <80 mL/min/1.73m^2^(17). The other study showed there were 19.3% of elderly over 60 years old whose GFR 60mL/min/1.73m^2^. Most of them combined with hypertension, diabetes, cardiovascular disease and renal impairment. Only 0.7% elderly had decreased GFR without other comorbidities and renal damage. Aging, hypertension and cardiovascular disease had correlation with decreased GFR(18). Since 70% morinidazole is excreted via kidney(3), the eGFR calculation is necessary for the elderly if they also present impaired liver function.

The limitation of this study is the small sample size of moderate liver function impaired subjects with mild renal function impairment. The other limitation is that a PK study with 500mg q24h regimen of morinidazole infusion among both liver and renal impaired population was not conducted to collect the AUC data and compare with AUC of 500mg q12h of morinidazole in healthy subject.

## Conclusion

The PPK model of morinidazole was validated by this study. Moderate liver function impairment and body weight were the significant covariates for CL and V, respectively. According to results of this study, it is not necessary for moderate hepatic impairment patients who are not combined with renal function impairment to adjust the morinidazole dosage. For patient with moderate hepatic and mild renal impairment, morinidazole administration should be considered as 500mg every 24 hours. If a patient has moderate/severe liver and renal function impaired, both dosage and interval adjustment of morinidazole should be considered. For patient with mild liver function and renal function impairment, an intensive safety monitoring of safety and therapeutic drug monitoring might be needed during the administration of morinidazole and dosage adjustment is required, if necessary. For elderly patients with liver function impairment, eGFR calculation is recommended before administration of morinidazole.

## Acknowledgments

Morinidazole is manufactured by Jiangsu Haosen Pharmaceutical Co. LTD. The PK study of morinidazole among moderate hepatic impairment patients is sponsored by Jiang Su Heng Rui Medicine Co. LTD in 2009 to 2010.

